# Approximate inference of gene regulatory network models from RNA-Seq time series data

**DOI:** 10.1101/149674

**Authors:** Thomas Thorne

## Abstract

Inference of gene regulatory network structures from RNA-Seq data is challenging due to the nature of the data, as measurements take the form of counts of reads mapped to a given gene. Here we present a model for RNA-Seq time series data that applies a negative binomial distribution for the observations, and uses sparse regression with a horseshoe prior to learn a dynamic Bayesian network of interactions between genes. We use a variational inference scheme to learn approximate posterior distributions for the model parameters. The methodology is benchmarked on synthetic data designed to replicate the distribution of real world RNA-Seq data. We compare our method to other sparse regression approaches and information theoretic methods. We demonstrate an application of our method to a publicly available human neuronal stem cell differentiation RNA-Seq time series.

## 1 Introduction

Methods for the inference of gene regulatory networks from RNA-Seq data are currently not as mature as those developed for microarray datasets. Normalised microarray data posses the desirable property of being approximately normally distributed so that they are readily amenable to various forms of inference, and in the literature many graphical modelling schemes have been explored that exploit the normality of the data[47, 18, 38, 24, 25, 15, 44, 45, 46].

The data generated by RNA-Seq studies on the other hand present a more challenging inference problem, as the data are no longer approximately normally distributed, and before normalisation take the form of non-negative integers. In the detection of differential expression in RNA-Seq data, negative binomial distributions have been applied [16, 3, 41, 26], providing a good fit to the over-dispersion typically seen in the data relative to a Poisson distribution. Following similar graphical modelling approaches as applied in the analysis of microarray data, it is natural to consider Poisson and negative binomially distributed graphical models. Unfortunately in many cases when applying graphical modelling approaches with Poisson distributed observations, only models that represent negative conditional dependencies are available, or inference is significantly complicated due to lack of conjugacy between distributions[19]. Poisson graphical models have been applied successfully in the analysis of miRNA regulatory interactions [2, 13], but we might expect to improve on these by modelling the overdispersion seen in typical RNA-Seq data sets with a negative binomial model.

One specific case of interest in the analysis of RNA-Seq data is the study of time series in a manner that takes into account the temporal relationships between data points. Previous work in the literature has developed sophisticated models for the inference of networks from microarray time series data [24, 25], but whilst methods have been developed for the analysis of differential behaviour in RNA-Seq time series [1, 21], little attention has been given to the task of learning networks from such data. Here we present a method for the inference of networks from RNA-Seq time series data through the application of a Dynamic Bayesian Network (DBN) approach, that models the RNA-Seq count data as being negative binomially distributed conditional on the expression levels of a set of predictors.

## 2 Methods

### 2.1 Dynamic Bayesian Networks

In a DBN framework[23], considering only edges between time points, we can model a sequence of observations using a first order Markov model, where the value of a variable at time *t* is dependent only on the values of a set of parent variables at time *t* − 1. This can be written as
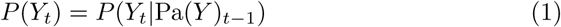

where Pa(*Y*) is the set of parents of variable *Y* in the network. In our case we wish to model the expression level of a gene conditional on a set of parent genes that have some influence on it. To learn the set of parent variables of a given gene, it is possible to perform variable selection in a Markov Chain Monte Carlo framework, proposing to add or remove genes to the parent set in a Metropolis-Hastings sampler. However this can be computationally expensive, and so instead we consider applying a sparse regression approach to learn a set of parents for each gene.

### 2.2 Sparse negative binomial regression

Given data *D* consisting of *M* columns and *L* rows, with columns corresponding to genes and rows to time points, we seek to learn a parent set for each gene. To do so we can employ a regularised regression approach that enforces sparsity of the regression coefficients, and only take predictors (genes) whose coefficients are significantly larger than zero as parents. To simplify the presentation, below we consider the regression of the counts for a single gene *i, y* = *D*_2*:L,i*_, conditional on the counts of the remaining *W* = *M* − 1 genes *X* = *D*_1:(*L*−1),−*i*_. The matrix *X* is supplemented with a column vector **1.** Where there are multiple replicates for each time point these can be adjusted appropriately.

The counts *y_t_* are then modelled as following a negative binomial distribution with mean exp (*Xβ*)*_t_* and dispersion *ω*, where *β* is a vector of regression coefficients. The model is then
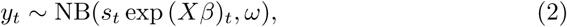

where we have applied the NB2 formulation of the negative binomial distribution, and *s_t_* is a scaling factor for each sample to account for sequencing depth. This corresponds to a DBN where
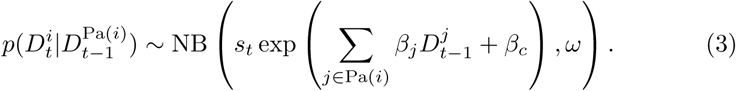

To enforce sparsity of the *β_w_* we apply a horseshoe prior[6, 7], assuming that 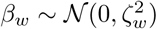, and placing a half-Cauchy prior on the 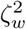,
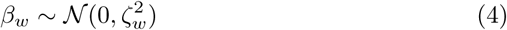

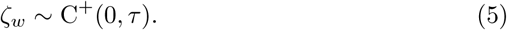

Then as in Carvalho et al. [7] we set a prior on *τ* that allows the degree of shrinkage to be learnt from the data
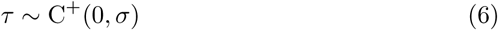

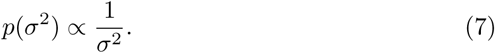

Finally we place a gamma prior on the dispersion parameter *ω.* This gives a joint probability
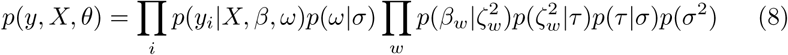

We now apply a variational inference[28, 29, 5, 4, 36] scheme to learn approximate posterior distributions over the model parameters. Unfortunately in this model the optimal distribution 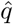 for the regression coefficients *β_w_* does not have a tractable solution. However following Luts and Wand [27] we can sidestep this problem by applying non-conjugate variational message passing[22], and we can then derive approximate posterior distributions for each of the model parameters following a straightforward parameter update scheme. The full set of variational updates are given in appendix A.

## 3 Results

### 3.1 Synthetic data

We apply our method to the task of inferring directed and undirected networks from 10 simulated gene expression time series. The time series were generated by utilising the GeneNetWeaver[43] software to first generate subnetworks representative of the structure of the *Saccharomyces cerevisiae* gene regulatory network, and then simulate the dynamics of the networks. Subnetworks of 50 nodes were generated and used to simulate 21 time points with 10 replicates, as in the DREAM4 challenge[39, 31, 32]. Simulated gene expression levels were then transformed by the inverse cumulative distribution function (CDF) *F*^−1^ of a normal distribution with mean and variance estimated from the data, and subsequently by the CDF of a negative binomial distribution *G.* The distributions *G* for each variable were chosen with mean and dispersion parameters randomly sampled from the empirically estimated means and dispersions of each gene from a publicly available RNA-seq count data set from the recount2 database[8] (accession *ERP003613)* consisting of 171 samples from 25 tissues in humans[10]. This was done so as to simulate the observed distributions of RNA-Seq counts in a real world data set.

We compare our approach against the Lasso as implemented in the lars R package[17] and the regularised Poisson regression method implemented in the glmnet R package[12]. The degree of regularisation was selected using cross validation as implemented in the respective software packages. For the undirected network inference case we also consider networks inferred by the mutual information based approaches implemented in the minet R package[35], specifically the CLR[11], ARACNE[33] and MRNET[34] methods.

In figure 1 we show the area under the receiver operating curve (AUC-ROC) and area under the precision recall curve (AUC-PR), as calculated by the PRROC R package[14], and Matthews Correlation Coefficient (MCC), for the various methods to be benchmarked in the undirected case. For the MCC, edges were predicted as those where zero was not contained in the 95% credible interval of the corresponding regression coefficients, and for the Lasso and glmnet methods, non-zero coefficients were taken as predicted edges. In the undirected case, we ignore the directionality of the edges in the true network generated by the GeneNetWeaver software, and take the strongest prediction in either direction to be the given score for an edge. When benchmarking the methods we found that there was variance in performance depending on the empirical means and dispersions of the negative binomial distributions that were sampled from the RNA-Seq data. Therefore we repeated benchmarking on each network 5 times with resampled negative binomial means and dispersions to take this into account. Running time for our algorithm was under 10 minutes for the 50 node networks considered.

**Figure 1:**
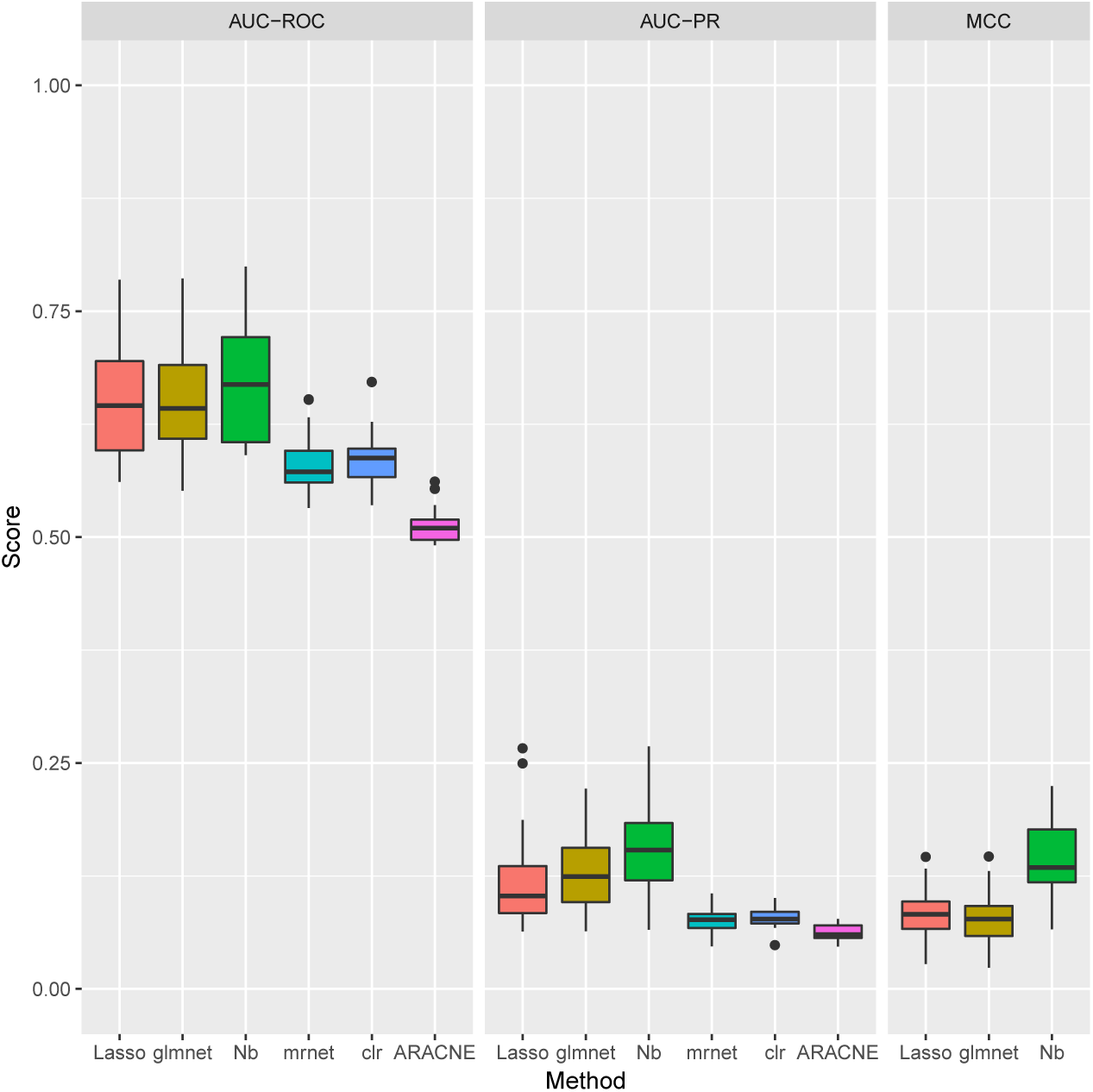
Boxplots of AUC-ROC, AUC-PR, and MCC for our method (Nb) and the competing methods benchmarked when learning undirected networks from synthetic data, for 10 subnetworks sampled from the *S. cerevisiae* gene regulatory network.

For the undirected case, our method shows an improved performance over the competing methods in terms of the AUC-PR, and also in terms of the MCC. Although the distinction between the approaches is less marked for the AUCROC, this is to be expected as the simulated biological network structures have far fewer (*<* 10%) true positives than true negatives, a situation in which the AUC-ROC does not distinguish performance as well as AUC-PR [9, 14].

As can be seen in figure 2 performance is slightly reduced in the directed case, likely due to the extra complexity of correctly predicting the directionality of edges. However our approach still clearly outperforms the competing methods in terms of the AUC-PR and MCC.

**Figure 2:**
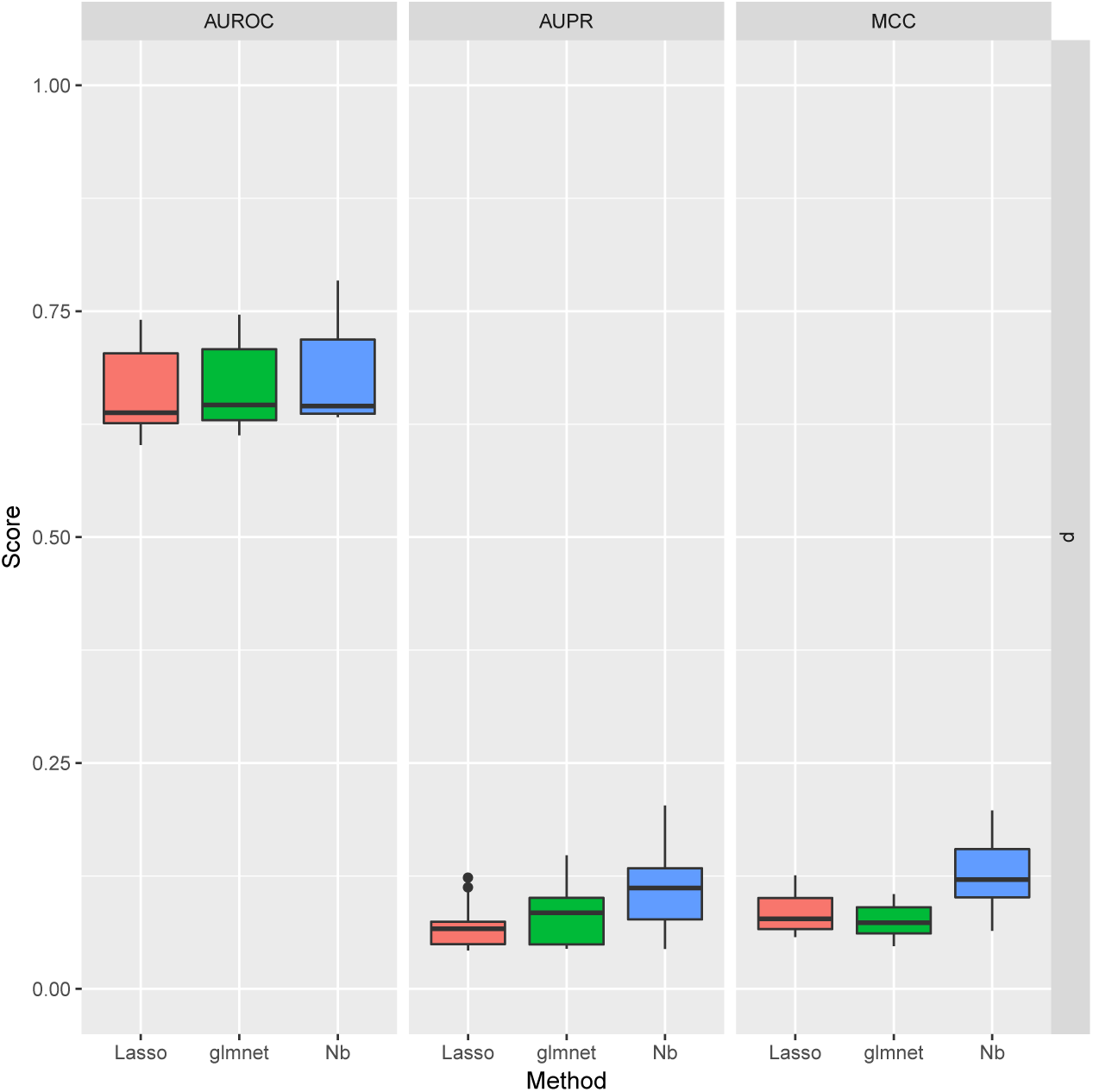
Boxplots of AUC-ROC, AUC-PR, and MCC for our method (Nb) and the methods benchmarked when learning directed networks from synthetic data, for 10 subnetworks sampled from the *S. cerevisiae* gene regulatory network.

It is clear that, as might be expected, taking into account the distributional properties observed in RNA-Seq data improves on the performance of methods based on assumptions that do not hold for RNA-Seq count data. We also see improvements over the nonparametric mutual information based approaches when learning undirected networks, perhaps due to our method taking into account the sequential nature of the data, and the constraint of assuming negative binomially distributed observations reducing the complexity of the inference problem.

## 4 Neural progenitor cell differentiation

To illustrate an application of our model to a real world RNA-Seq data set, we consider a publicly available RNA-Seq time course data set available from the recount2 database[8], accession *SRP041159.* The data consist of RNA-Seq counts from neuronal stem cells for 3 replicates over 7 time points after the induction of neuronal differentiation[42]. To select a subset of genes to analyse we performed a differential expression test between time points using the DESeq2 R package[26], and selected the 25 genes with the largest median fold-change between time points that were also differentially expressed between all time points. We chose to consider only an undirected network, as our benchmarking shows that such networks are likely more reliable.

Applying our method produced the network shown in figure 3, where it can be seen that there is a strongly interconnected network with four genes (MCUR1, PARP12, COL17A1, CDON) acting as hubs, suggesting these genes may be important in neuronal differentiation. Of these genes, CDON has been shown to be promote neuronal differentiation through the activation of p38MAPK pathway[48, 37] and inhibition of Wnt signalling[20], whilst MCUR1 is known to bind to MCU[30], that in turn has been shown to influence neuronal differentiation[40].

**Figure 3:**
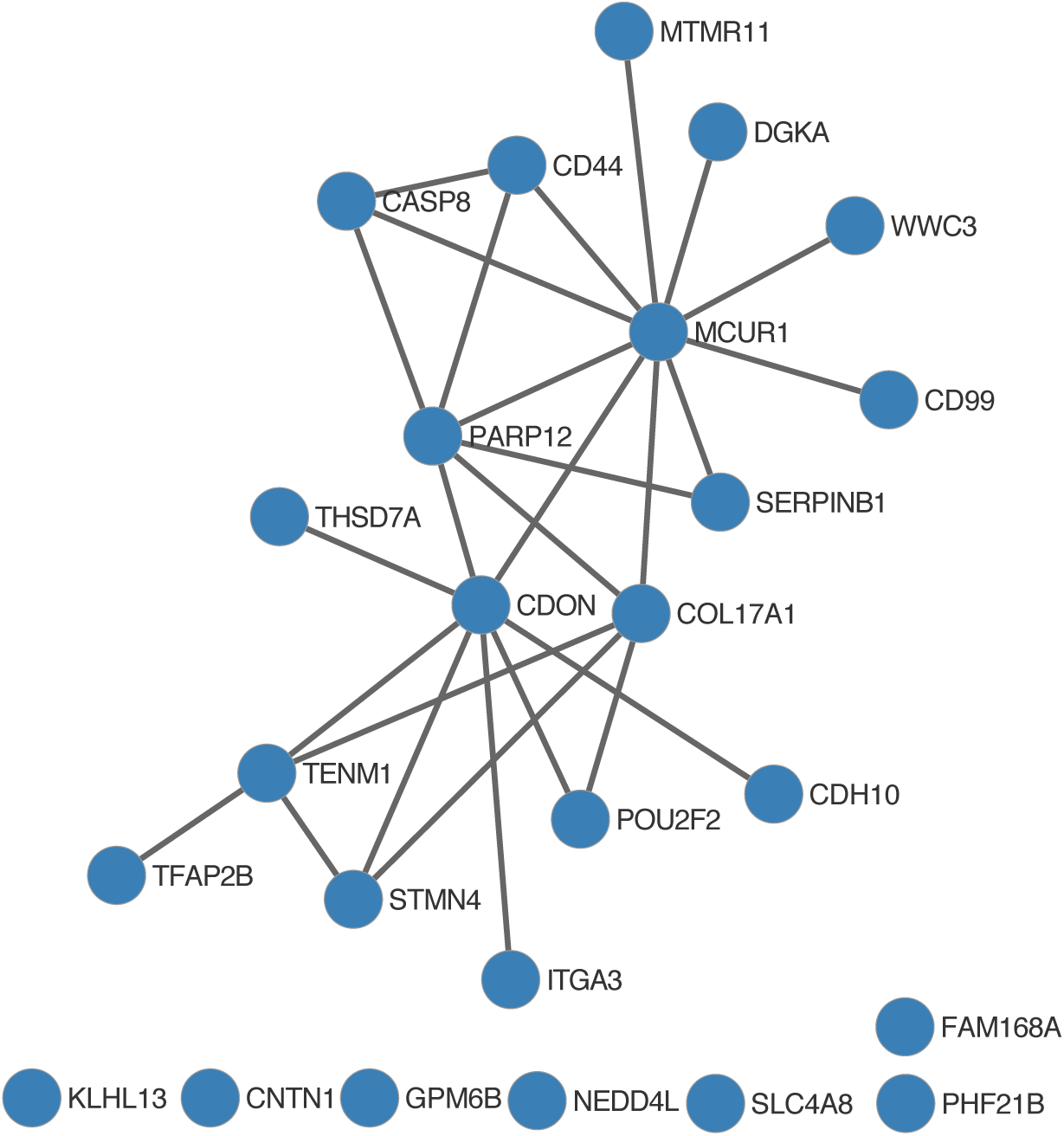
Undirected network inferred from the human neuronal differentiation time series data set.

## 5 Conclusions

We have developed a fast and robust methodology for the inference of gene regulatory networks from RNA-Seq data that specifically models the observed count data as being negative binomially distributed. Our approach outperforms other sparse regression and mutual information based methods in learning directed and undirected networks from time series data.

Another approach to network inference from RNA-Seq data could be to further develop mutual information based methodologies with this specific problem in mind. Mutual information based methods have the benefit of being independent of any specific model of the distribution of the data, and so could help sidestep issues in parametric modelling of RNA-Seq data. However this comes at the cost of abandoning the simplifying assumptions that are made by applying a parametric model that provides a reasonable fit to the data, and presents challenges of its own in the reliable estimation of the mutual information between random variables.

## Acknowledgement

T.T acknowledges the Edmond J. Safra Foundation and Lily Safra.

**A Variational inference**

From the results in Luts and Wand [27] the model can be written as a Poisson-Gamma mixture, so that
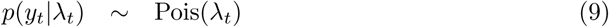

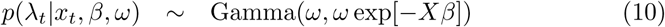

and the horseshoe prior on *β* represented using a mixture of inverse gamma distributions,
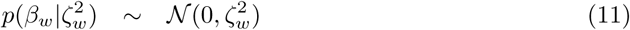

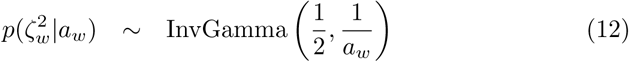

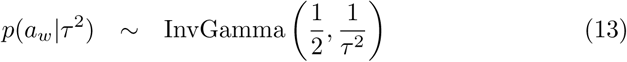

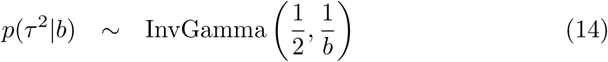

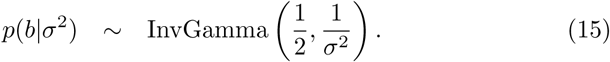

## A.1 Mean field approximation

The mean field approximation of the posterior is then
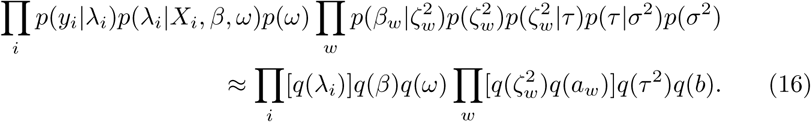

The variational updates for *λ_t_* can be derived as
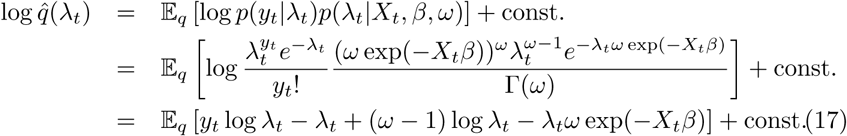

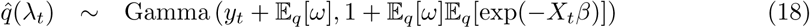

and due to the properties of the log-normal distribution
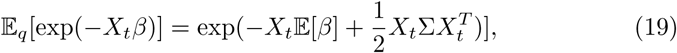

where Σ is the covariance matrix of *β* under 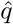.

As derived in Luts and Wand [27], applying non-conjugate variational message passing, 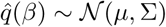 and the variational update for *β* is
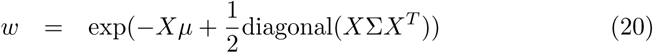

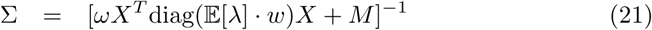

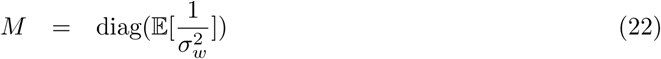

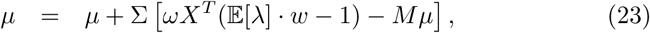

and for the dispersion *ω* we apply numerical integration as described in Luts and Wand [27].

Then for the horseshoe prior on *β*, the variational updates are
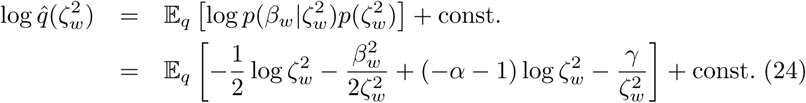

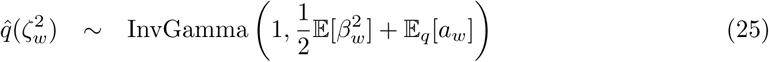

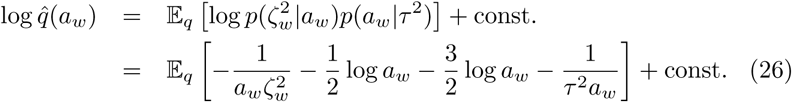

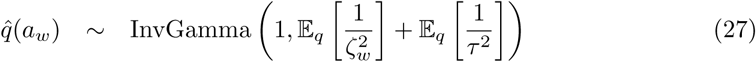

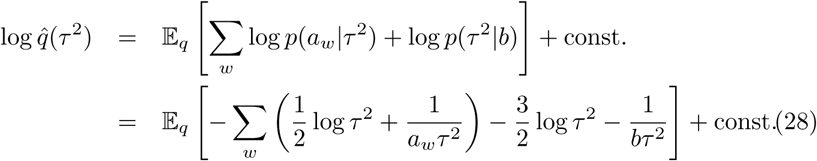

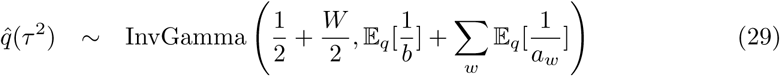

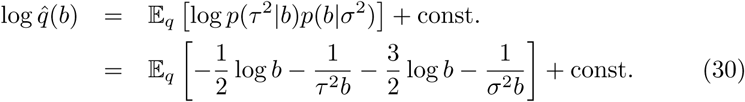

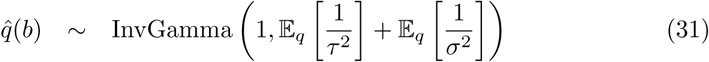

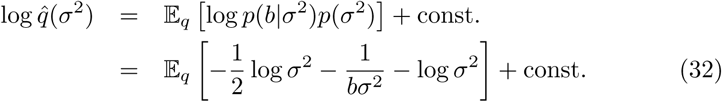

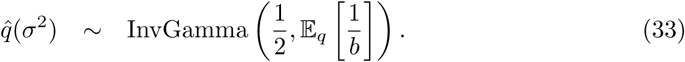

